# Redox potential in animal andrology: An experimental study on boar semen

**DOI:** 10.64898/2026.05.21.726780

**Authors:** Eliana Pintus, Maria Scaringi, Jef Engelen, José Luis Ros-Santaella

## Abstract

Impaired seminal redox balance is a main factor that contributes to male fertility disorders and reduced sperm survival during storage. Although several methods are available to measure antioxidant and reactive oxygen species (ROS) levels, their cost and complexity limit their use in routine sperm analysis. Recently, assessment of oxidation-reduction potential (ORP) has emerged as a convenient and comprehensive method for evaluating seminal redox status. While the implications of seminal ORP in humans have been extensively explored, its use in other species is limited. In this study, we explored the relationship between boar seminal ORP and sperm quality and its dynamics during liquid preservation. We found that the ORP of the porcine ejaculate was lower than that of the seminal plasma, while both parameters were correlated with the total antioxidant capacity (TAC) of seminal plasma. Sperm concentration and seminal pH influenced the seminal ORP, with lower values observed in ejaculates with higher sperm concentration and pH. Notably, a more oxidative seminal environment (characterized by high ORP or low TAC) was correlated with high mitochondrial activity and sperm velocity in fresh samples, which might be explained by increased ROS production by sperm mitochondria. Our results also show that seminal ORP increased during three days of liquid storage, while the ORP of the extender did not change significantly during the same period. Our findings advance our understanding of the implications of redox status in porcine sperm biology and pave the way for the broader application of ORP measurement in animal andrology.

**Highlights:**

- The ejaculate’s oxidation-reduction potential is lower than that of seminal plasma
- Seminal oxidation-reduction potential is correlated with total antioxidant capacity
- Seminal redox status is influenced by sperm concentration and semen pH
- An oxidative seminal environment is correlated with high sperm metabolism
- Seminal oxidation-reduction potential increases during 3 days of liquid storage

**Graphical abstract:** 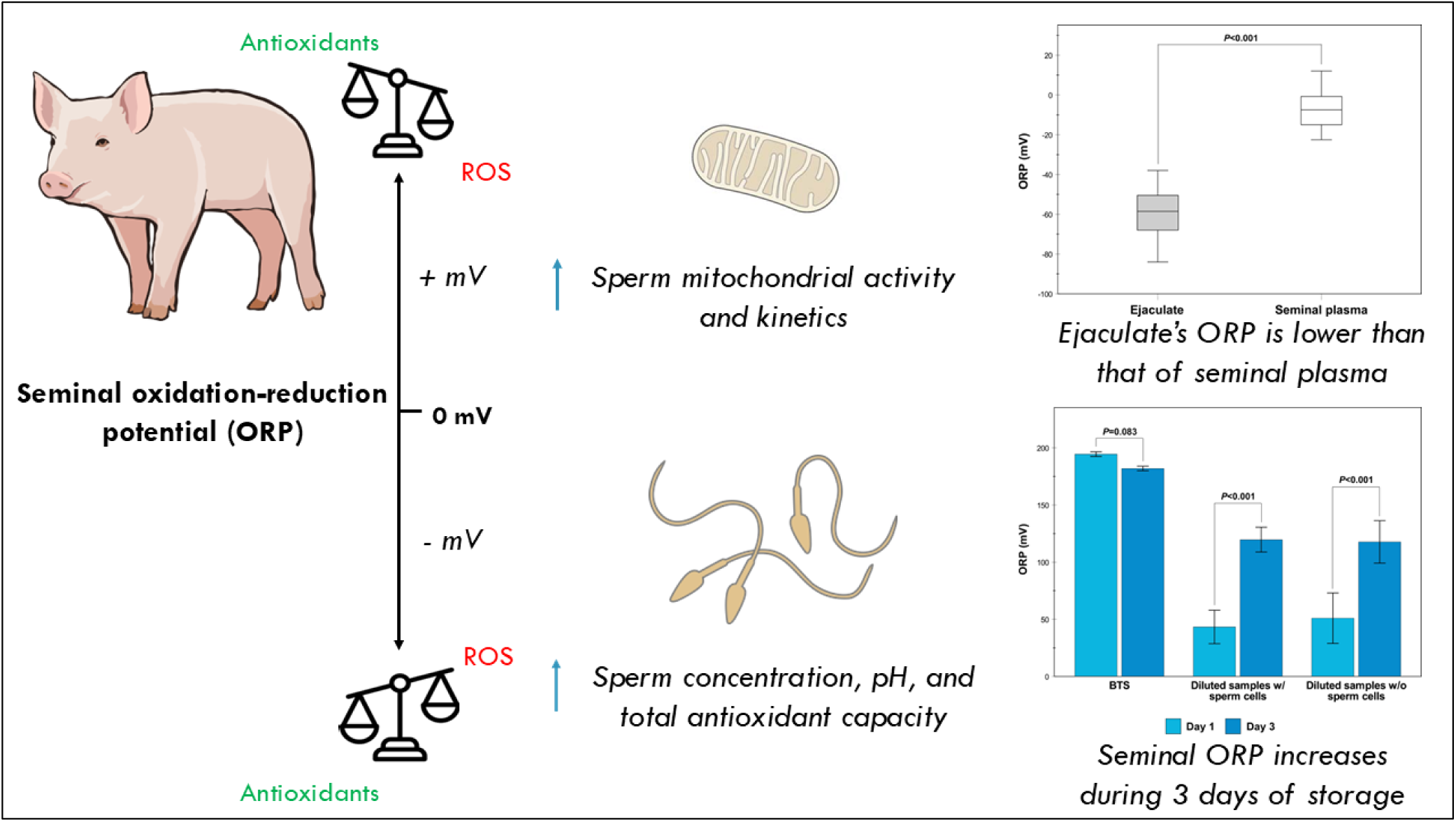

## 1. Introduction

Seminal redox status is defined by the balance between reactive oxygen species (ROS) and antioxidants in the ejaculate. An imbalance between these components can lead to either oxidative stress (an excess of ROS) or reductive stress (an excess of antioxidants). While oxidative stress has received more attention and is a well-established cause of male infertility in both humans [1] and other animals [2], recent research points out that reductive stress is equally harmful to sperm cells [3]. Due to the complexity of factors that contribute to sperm redox homeostasis, various methods have been developed to assess the levels of oxidants and antioxidants in the ejaculate [4]. However, quantifying these components – *in toto* or *ex parte* – as well as their detrimental consequences on cell function only provides a single dimension of seminal redox status.

In the last decade, the oxidation-reduction potential (ORP) has emerged as a rapid, simple, and effective method for the simultaneous evaluation of the ratio between seminal oxidants and antioxidants [4, 5]. By providing a comprehensive picture of the sperm redox status, ORP offers the potential to assess both oxidative and reductive stress in a biological system. It is important to note that ORP does not quantify the amount of antioxidants or oxidants; instead, it measures the electron transfer between molecules. The principle of ORP measurement is similar to that of pH, as both are measured by electrodes. However, while a pH electrode assesses the availability of protons in a solution, an ORP electrode reflects the potential flux of electrons [6]. The ORP values are expressed in millivolts (mV) and can range from negative to positive values. Negative values indicate a likely electron-donating, or reducing, environment, while positive values suggest a likely electron-accepting, or oxidative, environment. The measurement of ORP was originally applied in water disinfection [7] but has since been applied to biological fluids. For example, blood plasma ORP values are significantly increased in multi-trauma patients [8], those with head trauma [9], and individuals with metabolic syndrome and type II diabetes [10]. Elevated ORP has also been observed during surgical procedures [11], sepsis [12], and following excessive exercise [13].

In humans, seminal ORP has been correlated to testicular volume, hormone levels, sperm quality and freezability, and the outcomes of assisted reproductive technologies (ARTs), yet with controversial results ([14–18] but see also [19, 20]). It is important to acknowledge that seminal ORP values are often normalized for sperm concentration, hence expressed as mV/10^6^ sperm cells/mL. As a result, the intrinsic correlation between sperm concentration and other seminal parameters may have influenced the findings [6, 21]. Despite this, recent meta-analyses have shown that ORP measurement has high sensitivity and specificity for diagnosing male infertility and is positively associated with sperm DNA fragmentation [22, 23]. While further research is ongoing, seminal ORP measurement has been included as an advanced examination in the latest WHO laboratory manual for the examination and processing of human semen [24].

In contrast to its use in humans, the relevance of seminal ORP assessment in animal andrology is still in its early stages. In our previous study [25], we assessed porcine seminal ORP in samples exposed to hydrogen peroxide (H_2_O_2_) as a ROS-generating system, finding higher values in these treatments compared to control groups. In addition to H_2_O_2_, Balló et al. [26] found that seminal ORP values in humans were also increased by other ROS-generating systems, such as menadione or tert-butyl hydroperoxide. More recently [27], we found that nisin, a polypeptide bacteriocin, reduces the boar seminal ORP compared to treatments with or without antibiotic supplementation, providing evidence of antioxidant properties of this antimicrobial peptide. However, the physiological ORP value of boar ejaculate and its relationship with sperm quality and lifespan remain poorly understood [28]. In this study, we aimed to evaluate the ORP values in porcine ejaculate and seminal plasma, their relationship to the seminal antioxidant capacity and sperm parameters, and their dynamics during liquid storage of semen. The findings from this study advance our understanding of the implications of redox status in porcine sperm biology and pave the way for the broader application of ORP measurement in animal andrology.

## 2. Materials and Methods

### 2.1 Reagents

All reagents were purchased from Sigma-Aldrich (Prague, Czech Republic), unless otherwise stated.

### 2.2 Sample collection and experimental design

Twenty-seven ejaculates, each from a different and healthy Duroc boar, were collected from January 2023 to May 2023 at a pig breeding company (Lipra Pork, a.s., Czech Republic). Sperm-rich fractions of boar ejaculates were collected by the gloved-hand method and randomly split into two tubes, one of which was kept as raw semen, while another one was diluted 1:1 in Beltsville Thawing Solution (BTS), which was prepared as previously described [27]. After that, the samples were slowly cooled from 35 °C to 25 °C and transported within one hour to the laboratory. Here, raw and diluted semen were handled as described below:

#### Raw semen

Sperm concentration and morphology were assessed under phase-contrast microscopy (Eclipse E200, Nikon, Tokyo, Japan; 40× objective) after diluting an aliquot of each sample into phosphate-buffered saline (PBS) solution with 0.3% formaldehyde. Seminal plasma was obtained after double centrifugation at 2,000 *g* for 10 minutes at 17 °C. An aliquot of seminal plasma was used for the assessment of pH (Five Easy F20, Mettler-Toledo, Greifensee, Switzerland) and osmolality (Osmomat 3000, Gonotec, Berlin, Germany), with both analyses performed in duplicate. The remaining seminal plasma was either frozen at -80 °C for further analyses or diluted 1:3 into BTS to reach the final seminal plasma concentration of 25% and stored at 17 °C.

#### Diluted semen

Sperm samples were further diluted 1:1 with BTS to reach the seminal plasma concentration of 25% and slowly cooled from 25 °C to 17 °C. Once sperm concentration was established using a Bürker hemocytometer and under phase-contrast microscopy (Eclipse E200, Nikon, Tokyo, Japan; 40× objective), each ejaculate was diluted to 2×10^7^ spermatozoa/mL in BTS supplemented with 25% autologous seminal plasma. The resulting doses were stored at 17 °C for 3 days, in accordance with the established preservation limits for the BTS extender [29]. Both sperm and seminal plasma concentrations were standardized to mitigate the potential influence of these confounding variables. This specific concentration of seminal plasma was selected due to its documented beneficial effect on sperm quality during porcine semen storage [30]. Sperm motility, kinetics, mitochondrial activity, plasma membrane integrity, and acrosomal status were determined at day 1 and day 3 of semen storage after incubating the sample in a water bath at 38 °C for 20 minutes. At each incubation time, seminal ORP was determined with or without sperm cells, which were removed after centrifugation at 16,300 *g* for 5 minutes at room temperature. On day 1, an aliquot of each sample was also exposed to a ROS-generating system to assess lipid peroxidation levels.

### 2.3 Sperm analyses

#### 2.3.1 Seminal oxidation reduction potential (ORP)

The ORP was measured by a redox micro-electrode with Argenthal^TM^ reference system and platinum ring indicator (InLab Redox Micro, Mettler-Toledo, Greifensee, Switzerland) connected to a pH meter (Five Easy F20, Mettler-Toledo, Greifensee, Switzerland; resolution: 1 mV, accuracy: ± 1 mV), as previously described [25]. The ORP was recorded after embedding the microelectrode into a minimum of 0.7 mL of sample or BTS for 3 minutes at 38 °C. After each measurement, the probe was thoroughly washed with distilled water and calibrated into a redox buffer solution (220 mV, Mettler-Toledo, Greifensee, Switzerland) for 30 s. The assay was run in duplicate and expressed in millivolts (mV). The ORP value of the fresh boar ejaculate was also normalized for sperm concentration and expressed as mV/10^6^ sperm cells/mL [5, 31].

#### 2.3.2 Seminal total antioxidant capacity (TAC)

The TAC of the seminal plasma was determined by spectrophotometry, as previously described [32, 33]. The principle of this assay is based on the antioxidant’s capacity to reduce 2,2′-azinobis-(3-ethylbenzothiazoline-6-sulfonic acid) (ABTS), previously oxidized with H_2_O_2_. The TAC was measured on a 96-well plate by a multimode microplate reader (Biotek Synergy H1, Agilent, Santa Clara, CA, USA). The absorbance was measured at 660 nm immediately after adding a sample aliquot to reagent 1, and 5 minutes after adding reagent 2, as previously described [32]. A standard curve was established using known concentrations (31.25-2,000 µM) of 6-hydroxy-2,5,7,8-tetramethylchroman-2-carboxylic acid (Trolox), an analogue of vitamin E. The assay was run in duplicate for each sample and expressed as Trolox equivalents (μM). While several methods are available for measuring TAC, the ABTS decolorization assay was chosen because it is a popular method in andrology laboratories and is considered as an advanced technique for assessing oxidative stress, as specified in the WHO laboratory manual [24].

#### 2.3.3 Sperm mitochondrial activity, plasma membrane integrity, and acrosomal status

Sperm analyses were carried out as previously described [27]. Briefly, for the assessment of mitochondrial activity, sperm samples were incubated with rhodamine 123 (5 mg/mL in dimethyl sulfoxide, DMSO) and propidium iodide (0.5 mg/mL in PBS) for 15 minutes at 38 °C in the dark. Then, the samples were centrifuged at 500 *g* for 5 minutes. After discarding the supernatant, the spermatozoa were resuspended in PBS solution and evaluated using epifluorescence microscopy (Eclipse E600, Nikon, Tokyo, Japan; 40× objective). The sperm cells showing a bright green fluorescence over the midpiece were considered to have high mitochondrial activity. For the assessment of plasma membrane integrity, the sperm samples were incubated with propidium iodide (0.5 mg/mL in PBS), carboxyfluorescein diacetate (0.46 mg/mL in DMSO), and formaldehyde solution (0.3% in PBS) for 10 minutes at 38 °C in the dark. Then, the spermatozoa were assessed under epifluorescence microscopy (Eclipse E600, Nikon, Tokyo, Japan; 40× objective). The spermatozoa with green fluorescence over the entire head were considered to have an intact plasma membrane. For the acrosomal status, the sperm samples were fixed in a glutaraldehyde solution (2% in PBS) and evaluated under phase contrast microscopy (Eclipse E600, Nikon, Tokyo, Japan; 40× objective). The sperm cells with a normal apical ridge were considered to have an intact acrosome. Per each analysis, 200 sperm cells were evaluated by the same observer and counted using an app-based cell counter [34].

#### 2.3.4 Lipid peroxidation

Lipid peroxidation was induced by exposing sperm samples to the ROS-generating system composed of 50 µM FeSO_4_ and 500 µM sodium ascorbate during 3.5 hours at 38 °C [35]. At the end of the incubation, sperm samples were stored at -80 °C until analysis. Lipid peroxidation levels were assessed by the thiobarbituric acid reactive substances (TBARS) assay, as previously described [35]. Briefly, sperm samples were incubated into a solution 0.25 N HCl with 15% trichloroacetic acid and 0.375% thiobarbituric acid for 15 minutes at 90 °C. After that, the samples were transferred into ice-cold water for 5 minutes. Then, samples were centrifuged at 16,300 *g* for 5 minutes and the clear supernatant was transferred into a 96-well plate. The absorbance of each sample was measured at 532 nm by a multimode microplate reader (Biotek Synergy H1, Agilent, Santa Clara, CA, USA). A standard curve was established by using known concentrations of 1,1,3,3-tetramethoxypropane (0.5-32 µM), which, under acid hydrolysis, becomes malondialdehyde (MDA). The levels of lipid peroxidation are shown as μmol of MDA per 10^8^ spermatozoa. The assay was run in duplicate for each sample.

#### 2.3.5 Sperm motility and kinetics

Sperm motility and kinetics were assessed by a Computer Assisted Sperm Analyzer (CASA; NIS-Elements, Nikon, Tokyo, Japan and Laboratory Imaging, Prague, Czech Republic), after loading a sperm aliquot into a pre-warmed Leja chamber (Leja Products BV, Nieuw-Vennep, The Netherlands; chamber depth: 20 μm) [27]. Sperm motility parameters were determined on a minimum of 200 sperm cells per sample: total motility (TM, %), progressive motility (PM, %), average path velocity (VAP, µm s^-1^), curvilinear velocity (VCL, µm s^-1^), straight-line velocity (VSL, µm s^-1^), amplitude of lateral head displacement (ALH, μm), beat-cross frequency (BCF, Hz). The standard CASA settings were as follows: frames per second, 60; minimum of frames acquired, 31; number of fields analyzed, 6; VAP ≥ 10 µm s^-1^ to classify a spermatozoon as motile; straightness (i.e., VSL/VAP×100) ≥ 80% to classify a spermatozoon as progressive. All videos were visually checked by the same researcher to remove debris or erroneously crossed sperm tracks.

### 2.4 Statistical analyses

Data were analyzed with the statistical program SPSS, version 30 (IBM Inc., Chicago, IL, USA). The Shapiro–Wilk and Levene tests were used to check for normal distribution and homogeneity of variance, respectively. Based on the data distribution, paired-sample t-test or Wilcoxon signed-rank test were used to check for differences between data from two related samples (e.g., ORP from raw ejaculate and seminal plasma). Two-tailed Pearson or Spearman rank correlations were applied to assess the relationship between parameters for data that were normally or not normally distributed, respectively. Cluster analysis was performed for classifying samples based on their ORP or TAC values of their seminal plasma. These parameters were selected because both were measured within the seminal plasma fraction, ensuring consistent processing conditions. Firstly, the number of clusters was automatically determined by the two-step cluster analysis using the Euclidean distance measure and Schwarz’s Bayesian criterion. Then, the number of clusters obtained was used to set up the K-Means cluster analysis by using the iteration and classification method. An independent sample t-test or the Mann-Whitney U test was used to check for differences between males with “low” and “high” ORP or TAC values for data that were normally or not normally distributed, respectively. All analyses were performed on 27 boars, except for the ORP of the raw ejaculate that was assessed in 23 animals. Because each ejaculate was collected from a different boar, samples were treated as independent biological replicates. Data are shown as mean±SD. The statistical significance was set at *P* values <0.05.

## 3. Results

### 3.1 Assessment of porcine seminal redox status and its correlation with sperm parameters

Descriptive statistics of seminal redox status and sperm parameters in fresh porcine ejaculates are shown in Table 1. On average, the ORP of boar ejaculate was -58.41±12.88 mV (normalized ORP: -0.13±0.04 mV/10^6^ sperm cells/mL), while that of the seminal plasma was -7.44±8.75 mV. The ORP of the seminal plasma was remarkably higher than that of the ejaculate (*P*<0.001, Figure 1A), which suggests that the presence of the sperm cells affects the ORP values in boar semen. As shown in Table 2, sperm concentration was indeed significantly associated with the ORP of the ejaculate (*P*=0.008), but not with that of the seminal plasma (*P*>0.05). As expected, ORP values of ejaculate and seminal plasma were themselves correlated (r=0.822, *P*<0.001). Yet, TAC of the seminal plasma was also positively associated with sperm concentration (*P*<0.001, Table 2).

**Figure 1.**
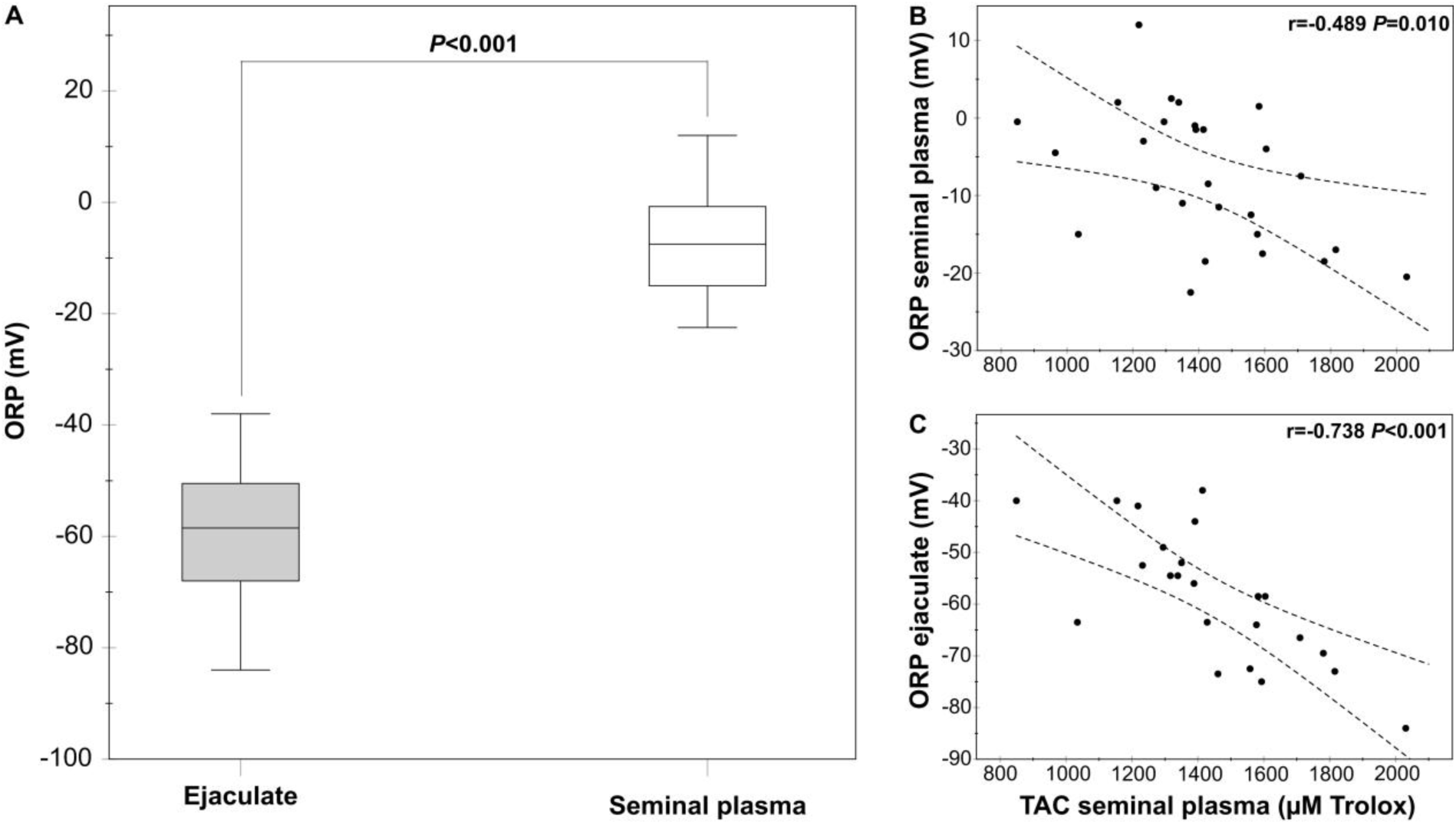
Redox status of fresh boar semen. A) Redox potential of ejaculate and seminal plasma. B) Correlation between redox potential and antioxidant capacity of seminal plasma. C) Correlation between ejaculate’s redox potential and seminal plasma’s antioxidant capacity. ORP: Oxidation-Reduction Potential; TAC: Total Antioxidant Capacity; *N*=23 for ejaculate ORP, *N*=27 for seminal plasma ORP and TAC.

**Table 1.** Descriptive statistics of seminal redox status and sperm parameters in porcine ejaculates.

|  | Mean±SD | Minimum | Maximum | 95% CI Lower | 95% CI Upper | N |
| --- | --- | --- | --- | --- | --- | --- |
| *Raw semen* |  |  |  |  |  |  |
| Ejaculate ORP (mV) | -58.41±12.88 | -84.00 | -38.00 | -63.98 | -52.85 | 23 |
| Normalized ORP (mV/10<sup>6</sup> sptz/mL) | -0.13±0.04 | -0.22 | -0.08 | -0.14 | -0.11 | 23 |
| Seminal plasma ORP (mV) | -7.44±8.75 | -22.50 | 12.00 | -10.91 | -3.98 | 27 |
| Seminal plasma TAC (μM Trolox) | 1,413.02±260.95 | 849.42 | 2,030.52 | 1,309.80 | 1,516.25 | 27 |
| Sperm concentration (10<sup>6</sup> sptz/mL) | 468.89±171.11 | 267.50 | 1,052.50 | 401.20 | 536.58 | 27 |
| pH | 7.48±0.18 | 7.29 | 8.05 | 7.41 | 7.55 | 27 |
| Osmolality (mOsm/Kg H<sub>2</sub>O) | 297.54±5.20 | 287.50 | 312.00 | 295.48 | 299.59 | 27 |
| Normal sperm morphology (%) | 63.11±16.76 | 27.00 | 87.50 | 56.48 | 69.74 | 27 |
| *<sup>§</sup>Diluted semen* |  |  |  |  |  |  |
| Intact plasma membrane (%) | 73.54±5.47 | 63.00 | 84.50 | 71.37 | 75.70 | 27 |
| Intact acrosome (%) | 95.44±1.75 | 91.00 | 98.50 | 94.75 | 96.14 | 27 |
| Lipid peroxidation (μM MDA/10<sup>8</sup> sptz) | 20.94±3.07 | 17.22 | 29.89 | 19.72 | 22.15 | 27 |
| Active mitochondria (%) | 77.00±5.86 | 61.50 | 85.00 | 74.68 | 79.32 | 27 |
| Total motility (%) | 76.22±11.06 | 47.02 | 92.25 | 71.84 | 80.59 | 27 |
| Progressive motility (%) | 58.32±13.26 | 21.33 | 81.79 | 53.07 | 63.57 | 27 |
| VAP (μm s<sup>-1</sup>) | 89.90±12.76 | 55.81 | 118.78 | 84.86 | 94.95 | 27 |
| VCL (μm s<sup>-1</sup>) | 140.80±24.59 | 68.57 | 181.33 | 131.08 | 150.53 | 27 |
| VSL (μm s<sup>-1</sup>) | 75.20±13.89 | 46.03 | 108.86 | 69.71 | 80.70 | 27 |
| ALH (μm) | 5.25±0.72 | 2.97 | 6.74 | 4.96 | 5.53 | 27 |
| BCF (Hz) | 16.83±1.86 | 13.44 | 20.62 | 16.10 | 17.57 | 27 |

<sup>§</sup>Samples were diluted to 2×10<sup>7</sup> sperm cells/mL into Beltsville Thawing Solution supplemented with 25% of autologous seminal plasma. ALH: Amplitude of Lateral Head Displacement; BCF: Beat-Cross Frequency; CI: Confidence Interval; MDA: Malondialdehyde; ORP: Oxidation-Reduction Potential; sptz: Spermatozoa; TAC: Total Antioxidant Capacity; VAP: Average Path Velocity; VCL: Curvilinear Velocity; VSL: Straight-Line Velocity.

**Table 2.** Correlations between porcine ejaculate redox status and sperm parameters.

|  | Ejaculate<br>ORP | Seminal<br>plasma ORP | Normalized<br>ORP | Seminal<br>plasma TAC |
| --- | --- | --- | --- | --- |
| *Raw semen* |  |  |  |  |
| Sperm concentration | -0.538** | -0.087 | 0.734*** | 0.625*** |
| pH | -0.466* | -0.516** | -0.080 | 0.186 |
| Osmolality | 0.285 | 0.189 | 0.134 | -0.324 |
| Normal sperm morphology | 0.134 | 0.090 | 0.141 | -0.233 |
| *<sup>§</sup>Diluted semen* |  |  |  |  |
| Intact plasma membrane | 0.107 | 0.042 | -0.220 | -0.221 |
| Intact acrosome | 0.199 | 0.247 | -0.339 | -0.338 |
| Lipid peroxidation | -0.423* | -0.553** | 0.187 | 0.293 |
| Active mitochondria | 0.585** | 0.292 | -0.248 | -0.491** |
| Total motility | 0.242 | 0.162 | -0.206 | -0.252 |
| Progressive motility | 0.118 | 0.111 | -0.236 | -0.440* |
| VAP | 0.467* | 0.037 | -0.016 | -0.426* |
| VCL | 0.141 | -0.021 | 0.095 | -0.081 |
| VSL | 0.494* | 0.109 | -0.140 | -0.551** |
| ALH | 0.294 | 0.024 | -0.014 | -0.339 |
| BCF | 0.186 | 0.004 | -0.005 | -0.205 |

<sup>§</sup>Samples were diluted to $2 \times 10^7$ sperm cells/mL into Beltsville Thawing Solution supplemented with 25% of autologous seminal plasma. ALH: Amplitude of Lateral Head Displacement; BCF: Beat-Cross Frequency; ORP: Oxidation-Reduction Potential; TAC: Total Antioxidant Capacity; VAP: Average Path Velocity; VCL: Curvilinear Velocity; VSL: Straight-Line Velocity. \* $P < 0.05$ ; \*\* $P < 0.01$ ; \*\*\* $P < 0.001$ ; $N = 23$ for ejaculate ORP and normalized ORP, $N = 27$ for seminal plasma ORP and TAC. Color intensity indicates the strength of correlation ranging from 1 (green) to -1 (blue).

High seminal TAC was significantly associated with low ORP value of boar ejaculate and seminal plasma (*P*≤0.01, Figures 1B and 1C), but not with normalized ORP value of the ejaculate (*P*>0.05). Moreover, low ORP values of both ejaculate and seminal plasma were associated with more alkaline pH (*P*=0.025 and *P*=0.006, respectively, Table 2). As shown in Table 2, ejaculates with low ORP or high TAC showed low mitochondrial activity, VAP, and VSL (*P*<0.05). Low seminal TAC was also associated with high progressive motility (*P*=0.022). Moreover, low ORP values of both ejaculate and seminal plasma were significantly correlated to high levels of lipid peroxidation (*P*=0.044 and *P*=0.003, respectively). Neither the ORP of raw ejaculate or seminal plasma was associated with seminal osmolality, percentage of morphologically normal spermatozoa, acrosomal status or plasma membrane integrity. The normalized ORP of the boar semen was significantly correlated with sperm concentration (*P*<0.001), but not with any other sperm parameter (*P*>0.05).

### 3.2 Use of porcine seminal redox status as a predictor of sperm quality during liquid storage

The boar ejaculates were classified into different groups based on their seminal plasma ORP or TAC in fresh samples. Using ORP of seminal plasma as the input variable, cluster analyses (incorporating both two-step and K-Means methods) identified two groups with similar sample size but different ORP values (*P*<0.001): a low SP-ORP cluster (-14.61±4.71 mV, *n*=14) and a high SP-ORP cluster (0.27±4.19 mV, *n*=13). As shown in Figure 2 and Table 3, ejaculates with low SP-ORP value showed significantly higher pH and lipid peroxidation levels than those with high SP-ORP (*P*<0.05) at day 1 of semen storage. Albeit not significant, cluster including ejaculates with low SP-ORP values also showed lower percentage of normal sperm, VAP, and VSL compared to those with high SP-ORP (*P≥*0.05, Figure 2 and Table 3). There were no differences between groups in other sperm parameters (*P*>0.05, Figure 2 and Table 3).

**Figure 2.**
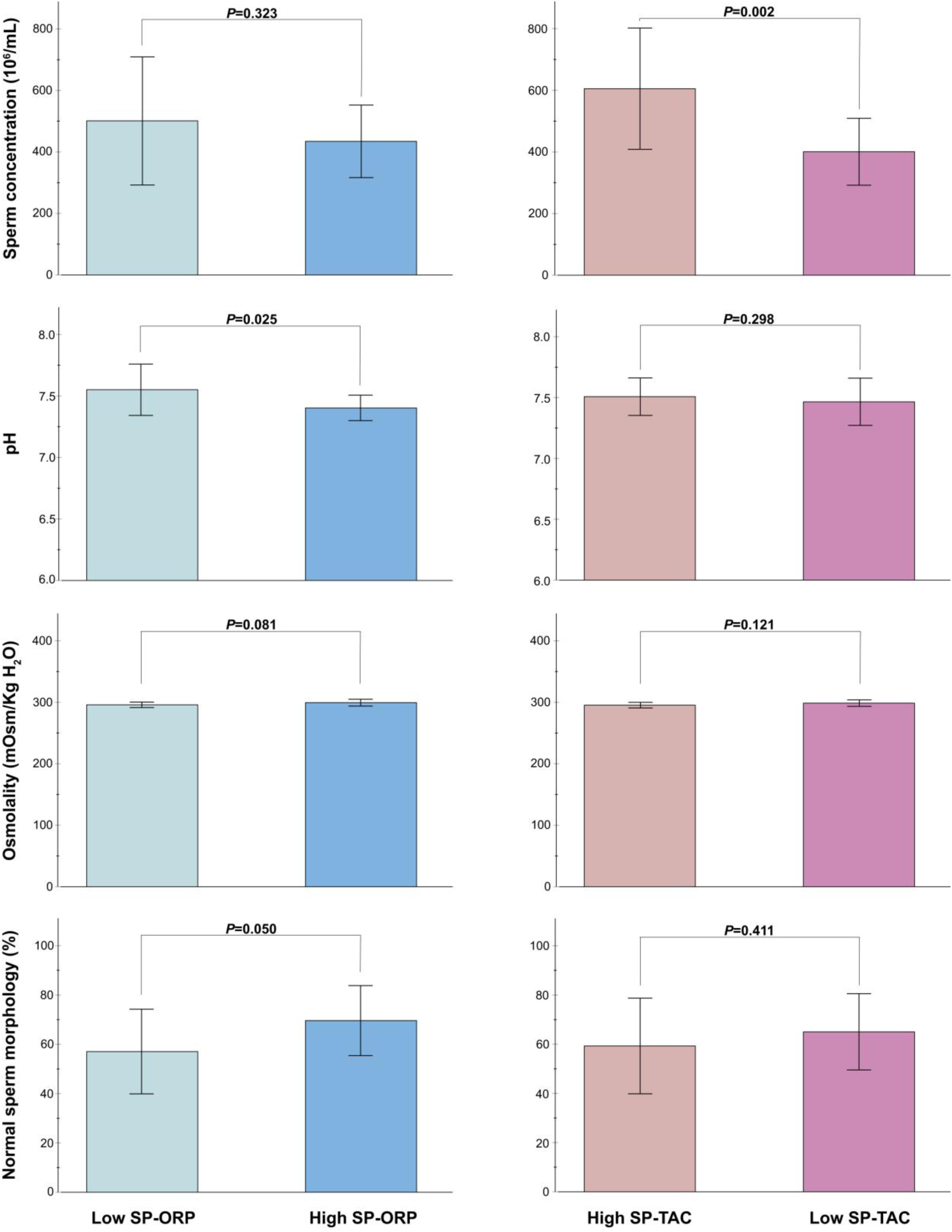
Cluster comparison of fresh seminal parameters in boar ejaculates with divergent redox profiles. ORP: Oxidation-Reduction Potential; SP: Seminal Plasma; TAC: Total Antioxidant Capacity; *N*=27.

**Table 3.** Cluster comparison of boar ejaculates with different redox status during three days of storage at 17 °C.

|  | Low SP-ORP<br>(n=14) | High SP-ORP<br>(n=13) | *P* | High SP-TAC<br>(n=9) | Low SP-TAC<br>(n=18) | *P* |
| --- | --- | --- | --- | --- | --- | --- |
| *Day 1* |  |  |  |  |  |  |
| Intact plasma membrane (%) | 72.96±5.38 | 74.15±5.72 | 0.583 | 72.28±5.35 | 74.17±5.58 | 0.409 |
| Intact acrosome (%) | 95.21±2.22 | 95.69±1.07 | 0.479 | 95.00±2.57 | 95.67±1.18 | 0.477 |
| Lipid peroxidation (μM MDA/10<sup>8</sup> spz) | 22.16±3.50 | 19.62±1.88 | **0.022** | 21.41±2.97 | 20.70±3.18 | 0.298 |
| Active mitochondria (%) | 75.43±6.98 | 78.69±3.93 | 0.151 | 73.11±5.34 | 78.94±5.20 | **0.012** |
| Total motility (%) | 73.40±12.54 | 79.25±8.69 | 0.155 | 71.07±14.24 | 78.79±8.42 | 0.087 |
| Progressive motility (%) | 56.07±14.82 | 60.74±11.44 | 0.372 | 52.53±15.62 | 61.21±11.30 | 0.110 |
| VAP (μm s<sup>-1</sup>) | 85.41±13.82 | 94.75±9.83 | 0.056 | 84.65±7.14 | 92.53±14.25 | 0.133 |
| VCL (μm s<sup>-1</sup>) | 137.98±30.72 | 143.84±16.37 | 0.547 | 136.92±23.88 | 142.74±25.38 | 0.275 |
| VSL (μm s<sup>-1</sup>) | 70.36±14.78 | 80.42±11.16 | 0.058 | 68.91±10.89 | 78.35±14.42 | 0.097 |
| ALH (μm) | 5.08±0.87 | 5.43±0.47 | 0.213 | 5.06±0.46 | 5.34±0.82 | 0.360 |
| BCF (Hz) | 16.48±1.98 | 17.22±1.72 | 0.155 | 16.30±1.94 | 17.10±1.82 | 0.298 |
| *Day 3* |  |  |  |  |  |  |
| Intact plasma membrane (%) | 65.71±6.41 | 71.35±5.01 | **0.018** | 66.33±6.67 | 69.47±6.10 | 0.233 |
| Intact acrosome (%) | 93.07±2.82 | 93.19±2.79 | 0.912 | 92.28±3.13 | 93.56±2.53 | 0.263 |
| Active mitochondria (%) | 71.71±6.20 | 72.19±7.26 | 0.855 | 69.94±4.66 | 72.94±7.30 | 0.274 |
| Total motility (%) | 56.58±20.12 | 58.10±14.34 | 0.825 | 57.09±16.95 | 57.42±17.90 | 0.965 |
| Progressive motility (%) | 44.40±8.97 | 51.77±10.30 | 0.058 | 44.73±9.37 | 49.56±10.42 | 0.253 |
| VAP (μm s<sup>-1</sup>) | 77.51±20.30 | 83.13±20.78 | 0.484 | 79.06±20.91 | 80.79±20.63 | 0.839 |
| VCL (μm s<sup>-1</sup>) | 125.85±42.88 | 122.74±37.16 | 0.843 | 126.19±43.68 | 123.43±38.52 | 0.868 |
| VSL (μm s<sup>-1</sup>) | 61.49±13.87 | 70.72±17.76 | 0.143 | 63.36±15.24 | 67.22±17.00 | 0.571 |
| ALH (μm) | 4.71±1.35 | 4.86±1.31 | 0.772 | 4.93±1.49 | 4.71±1.25 | 0.705 |
| BCF (Hz) | 16.43±1.99 | 17.40±2.08 | 0.226 | 16.28±2.38 | 17.20±1.87 | 0.278 |

Samples were diluted to 2×10<sup>7</sup> sperm cells/mL into Beltsville Thawing Solution supplemented with 25% of autologous seminal plasma. ALH: Amplitude of Lateral Head Displacement; BCF: Beat-Cross Frequency; MDA: malondialdehyde; ORP: Oxidation-Reduction Potential; SP: Seminal Plasma; TAC: Total Antioxidant Capacity; VAP: Average Path Velocity; VCL: Curvilinear Velocity; VSL: Straight-Line Velocity. Data are shown as mean±SD.

While the two-step cluster analysis initially suggested a single cluster for TAC, samples were subsequently classified into two groups based on a predefined number of clusters to maintain consistency with the ORP grouping variables. K-Means cluster analysis yielded two groups that showed significantly different TAC levels (*P*<0.001): a high SP-TAC cluster (1,694.65±157.45 µM Trolox equivalents, *n*=9) and a low SP-TAC cluster (1,272.21±171.73 µM Trolox equivalents, *n*=18). On day 1 of semen storage, boars with high SP-TAC in their seminal plasma showed significantly higher sperm concentration in their ejaculate and lower mitochondrial activity compared to those with low SP-TAC (*P*<0.05, Figure 2 and Table 3). Similarly to differences found between ORP clusters, samples with high SP-TAC also showed lower percentage of normal sperm cells, VAP, and VSL, although differences were not statistically significant (*P*>0.05, Figure 2 and Table 3).

Cluster analyses of porcine ejaculates, based on their ORP or TAC values in seminal plasma, did not reveal any further differences between groups after 3 days of liquid storage (Table 3). The sole exception observed was a higher percentage of sperm cells with intact plasma membrane in cluster with high SP-ORP cluster compared to the low SP-ORP cluster (*P*=0.018). Although differences were not statistically significant, the high SP-ORP cluster also showed high progressive motility and VSL compared to the low SP-ORP group (*P*>0.05, Table 3).

### 3.3 Effect of liquid storage on porcine seminal ORP with standardized seminal plasma and sperm concentration

The ORP values were still influenced by the presence of sperm cells at day 1 (43.28±14.63 mV with sperm cells *vs.* 50.96±22.09 mV without sperm cells, *P*<0.001), but not at day 3 (119.67±10.90 mV with sperm cells *vs.* 117.74±18.60 mV without sperm cells, *P*=0.639). Irrespective of sperm presence, ORP values significantly increased during 3 days of semen storage (Figure 3A). On the other hand, ORP of BTS slightly decreased from day 1 to day 3 of storage (194.50±2.00 mV *vs*. 182.00±2.00 mV, respectively), although differences were not statistically significant (*P>*0.05, *N*=3, Figure 3A). As shown in Figure 3B and 3C, the ORP values during semen storage followed a consistent pattern across clusters based on the TAC or ORP values of fresh seminal plasma. On day 1, samples with high SP-TAC exhibited lower ORP values compared to those with low SP-TAC, regardless of the presence of sperm cells (*P*<0.01, in samples with or without sperm cells, Figure 3C). On day 3, samples with high SP-TAC still showed lower ORP values than those with low SP-TAC, although differences were not statistically significant (*P*>0.05, Figure 3C).

**Figure 3.**
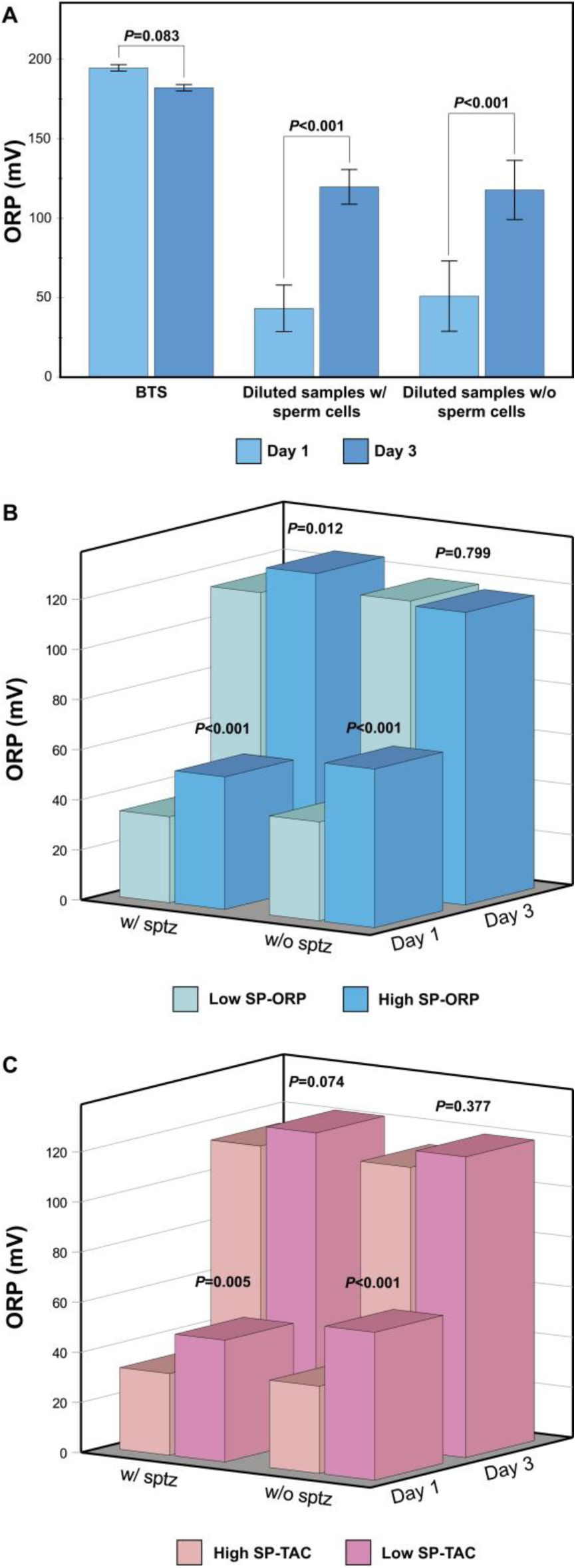
Dynamics and cluster comparison of seminal redox potential in boar ejaculates with divergent redox profile during 3 days of storage at 17 °C. A) Effect of liquid storage and sperm cells on seminal oxidation-reduction potential; B) Cluster comparison of redox potential dynamics during liquid storage in fresh ejaculates with low or high oxidation-reduction potential; C) Cluster comparison of redox potential dynamics during liquid storage in fresh ejaculates with low or high total antioxidant capacity. Samples were diluted to 2×10^7^ sperm cells/mL into Beltsville Thawing Solution supplemented with 25% of autologous seminal plasma. BTS: Beltsville Thawing Solution; ORP: Oxidation-Reduction Potential; SP: Seminal Plasma; Sptz: Spermatozoa; TAC: Total Antioxidant Capacity; w/: with; w/o: without; *N*=27.

## 4. Discussion

Our findings reveal that the ORP of the boar ejaculate and seminal plasma were significantly correlated with their TAC. To the best of our knowledge, this is the first time that a correlation between seminal ORP and TAC has been found. Our results also show that sperm concentration influenced the ORP of the ejaculate, while not that of the seminal plasma. On the other hand, the ORP of both ejaculate and seminal plasma were influenced by pH, with more alkaline conditions corresponding to lower ORP values. In contrast, while the TAC of seminal plasma also increased with sperm concentration, it was not affected by seminal pH. Unexpectedly, a more reductive seminal environment (characterized by low ORP or high TAC) was correlated with decreased mitochondrial activity and sperm velocity in fresh samples. This finding might be attributable to reduced levels of ROS produced by sperm mitochondrial metabolism. Our results also show that seminal ORP increased during three days of liquid storage irrespective of the presence of sperm cells. Taken together, our data provide the first evidence of a strong correlation between seminal ORP and TAC in a mammalian species. However, despite their similar relationships with sperm parameters, neither ORP nor TAC appeared to predict sperm quality during a three-day liquid storage period.

### 4.1 Redox status of fresh boar semen

On average, the absolute ORP value of the boar ejaculate was ∼-58 mV, which is remarkably lower compared to the ones of ∼45 mV reported in the human ejaculate [36, 37]. Given the influence of sperm concentration on ORP values found in our study, it is reasonable to hypothesize that the difference in ORP values between these species can be at least partly explained by the remarkable variation in their sperm concentration (∼469×10^6^ spermatozoa/mL in this study *vs*. ∼97×10^6^ spermatozoa/mL in the study by Joao et al. [37]). In line with our findings, Gill et al. [38] found that both absolute and normalized ORP values of human semen were negatively correlated with sperm concentration, that is, the lower the ORP value, the higher the sperm concentration. However, other studies in the same species did not find evidence of correlation between absolute ORP values and sperm number [20, 31, 39]. The remarkable difference in sperm concentration between swine and human species may also contribute to explain why the ORP values do not change between ejaculate and seminal plasma in humans [37], unlike in pigs. Our results indeed show that the ORP of porcine ejaculate is significantly lower compared to that of the seminal plasma. Another non mutually exclusive explanation for different ORP values between human and porcine ejaculate might concern methodological aspects, as seminal ORP in humans has been mostly assessed by the MiOXSYS^®^ device, which has been available on the market since 2016 [5]. While the principle at the basis of our and MiOXSYS^®^ devices are similar, they show some differences: i) in the MiOXSYS^®^ system, the final ORP reading is the average of the 20 readings that are automatically shown on the display screen at the end of the incubation period of approximately 2 minutes [36], while in our system ORP is constantly monitored during 3 minutes of measurement; ii) in the MiOXSYS^®^ device, sample volume is considerably lower (30 μL; [5]) than that required in our settings (i.e., ≥ 0.7 mL); iii) in the MiOXSYS^®^ system, samples are measured at room temperature [5], while in our experimental design ORP was measured at 38 °C. Because temperature is a key variable in the Nernst equation used to calculate ORP [4, 40] and has been shown to influence seminal ORP values [36], seminal redox status must be measured at a constant, controlled temperature to ensure accuracy and reproducibility. On the other hand, we found that the TAC of porcine seminal plasma was on average ∼1.4 mM Trolox equivalents, a value that is similar to the 1.8 mM described in fertile men [41] but higher compared to a previous study in boars where it ranged from 0.31 to 0.96 mM [42]. While the methodological procedures used in our and their study are similar, several factors, like individual variability or ejaculate fractions, may influence the TAC of boar seminal plasma [42]. Our findings show that boar seminal ORP and TAC are strongly correlated, providing further evidence that the former is a useful parameter for the assessment of the redox balance in the ejaculate.

### 4.2 Relationships between absolute or normalized seminal ORP and sperm traits

Several studies explored the relationship between sperm parameters and seminal ORP [5, 14, 15, 18, 19], often finding a significant correlation, especially when the latter is normalized for sperm concentration. This might be due to the fact that the normalized ORP value can lead to spurious correlations given the intrinsic relationship between sperm concentration and seminal parameters [6, 21]. In humans, for instance, normalized ORP values are negatively associated with sperm concentration, motility, and normal morphology [14, 43]. In contrast, absolute ORP values do not show any clear and consistent relationship with semen parameters [20]. Similarly, Joao et al. [37] and Castleton et al. [39] found a correlation between normalized ORP values and several sperm parameters (e.g., motility, morphology, DNA fragmentation), most of which, however, disappeared while considering the absolute ORP values. In another study in humans, Gill et al. [38] found that both absolute and normalized ORP values were correlated with sperm concentration, motility, morphology, and viability. Because of these inconsistencies, our study aimed to explore the relations between seminal redox balance and sperm parameters using both the absolute and normalized ORP values. Our findings show that absolute ORP values of boar ejaculate were associated with several sperm parameters (i.e., concentration, pH, velocity, mitochondrial activity, and lipid peroxidation), while only a significant correlation with sperm concentration was found with normalized ORP values. In contrast to previous findings in humans [14, 19, 38], there were no significant correlations between absolute or normalized ORP values and sperm morphology. This is noteworthy because the boars in this study were not pre-selected based on their spermiogram and exhibited significant variability in their seminal parameters. While the ejaculate parameters remained within the normal range described for this species [44], the percentage of normal sperm morphology was below typical averages. This result may be attributed to the highly stringent morphological criteria employed in our laboratory, as well as breed-specific characteristics previously documented in the Duroc population in the Czech Republic [45]. Collectively, our findings suggest that the relationship between seminal redox status and sperm parameters may vary among species and highlight the importance of using both absolute and normalized ORP values in andrology studies. While absolute values are particularly useful for descriptive or correlational studies, normalized values should be preferably considered in trials that investigate the effects of antioxidants or pro-oxidants on sperm function due to the influence of sperm concentration on seminal redox potential. In addition to sperm concentration, our findings also show that ORP of both boar ejaculate and seminal plasma were negatively correlated with the seminal pH. While no evidence has been found in human studies [19, 37, 46], it is known in other solutions that ORP decreases in alkaline pH [40]. The discrepancy between studies may be related to methodological differences. These include, but are not limited to, the use of pH meters *versus* pH strips and the reporting of normalized *versus* absolute ORP values. Using pH strips, Garcia-Segura et al. [46] did not find any correlation between normalized ORP and pH values in human semen, although the authors found that measurements of redox status became more variable at pH higher than 8. In our study, boar seminal pH was generally below 8 with 95% confidence interval between 7.41 and 7.55. In agreement with our findings, comparison of ORP in media used in human *in vitro* fertilization and andrology laboratories has shown that the lowest ORP values are found in the media with the most alkaline pH [47].

### 4.3 High sperm velocity and mitochondrial activity correlate with a more oxidative seminal environment

Interestingly, we also found that an oxidative seminal environment, either characterized by low TAC or high ORP values, was associated with high sperm velocity (specifically, VAP and VSL) and mitochondrial activity. This pattern remained consistent when comparing clusters with oxidative profiles (high SP-ORP or low SP-TAC) against those exhibiting a more reductive environment (low SP-ORP or high SP-TAC) on day one of semen storage. Since sperm mitochondria are a major source of ROS production [2] and boar sperm motility and velocity heavily rely on mitochondrial oxidative phosphorylation [48], intense sperm metabolism may likely contribute to this oxidative environment. In stallions, for instance, Gibb et al. [49] found that lipid peroxidation and ROS levels were positively associated with sperm motility and kinetics. In the same study, the authors also found that seminal oxidative stress was associated with improved pregnancy rates. Consequently, future research should focus on elucidating the specific mechanisms linking seminal redox status to sperm metabolic pathways, including oxygen consumption rates, ATP levels, and ROS generation. In agreement with Furtado et al. [20], we also found that the ORP values of the ejaculate were negatively correlated with lipid peroxidation. This relationship was further supported by comparing clusters with high *versus* low ORP values in their seminal plasma. Conversely, Castleton et al. [39] found a positive association between seminal ORP and lipid peroxidation in humans. It is remarkable to highlight that, in our study, lipid peroxidation was induced by exposing sperm samples to Fe^2+^/Ascorbate as a ROS-generating system with standardized sperm and seminal plasma concentration. In such ROS-generating system, a reducing agent like ascorbate can exhibit pro-oxidant effects by recycling Fe^3+^ back to Fe^2+^, thus catalyzing the formation of ROS [50, 51]. Therefore, lower ORP values might indicate greater antioxidant activity from specific seminal plasma compounds that could, in turn, boost the potency of the ROS-generating system, leading to a prooxidant effect. Further studies are required to test this hypothesis by assessing the redox potential of different ROS-generating systems.

### 4.4 Effect of liquid storage on seminal redox status

Our results show that seminal ORP significantly increased during a three-day period of liquid storage, independent of the presence of sperm cells. By the third day of storage, ORP values reached approximately a two-fold increase relative to initial measurement on day one, which indicates that seminal redox status undergoes a systemic convergence toward a more oxidative state over time. A comparative analysis of ejaculate clusters revealed on day one of semen storage that those characterized by a more oxidative profile (high SP-ORP or low SP-TAC) differed in sperm concentration, seminal pH, lipid peroxidation levels, mitochondrial activity, and redox status from those exhibiting a more reductive environment (low SP-ORP or high SP-TAC). However, these initial differences largely disappeared by day three of storage, which overall suggest that cluster classification of boar seminal samples based on their TAC or ORP in fresh seminal samples seem not to foresee sperm quality during 3 days of storage at 17 °C. In agreement with our findings, Barranco et al. [42] did not find that boar seminal TAC predicts sperm quality during the same storage period and conditions, although greater sperm decline was found in ejaculates with low seminal TAC. While human seminal ORP does not change over 4 hours post-ejaculation [36], Saleh et al. [16] found in the same species that seminal ORP increases approximately threefold after sperm cryopreservation compared to pre-freezing values. Whether the increase in ORP values during storage found in our study is due to increased ROS levels, decreased antioxidant protection, or both factors, remains to be investigated. Moreover, the increased ORP in freshly diluted semen (∼43 mV) compared to the raw ejaculate (∼-58 mV) may be attributed to several factors: i) addition of BTS extender (which exhibits a high baseline ORP of ∼194.5 mV); ii) dilution of seminal antioxidants and sperm cells; iii) and potential increased sperm ROS production resulting from mechanical handling of seminal doses. Another factor that may contribute to the increase of ORP values during storage is the composition of the extender used for semen preservation. In stallions, seminal extenders contain a considerable amount of glucose that exceeds the physiological range of this carbohydrate in the seminal plasma (67 *vs*. 5 mM) and might be responsible for glucose-induced oxidative stress [52]. Remarkably, the physiological concentration of glucose in boar seminal plasma is below 0.3 mM [53], while that of the BTS is 205 mM [54]. Regarding this aspect, our results also show that while the ORP of seminal doses increased during 3 days of semen storage, the ORP of BTS did not significantly change during this storage period. We also found that the ORP of BTS was about ∼194.5 mV, which is lower compared to the values reported by Panner Selvam et al. [47] in human ART media, in which ORP ranged from 209 to 274 mV. The assessment of the ORP of sperm washing and storage media is essential for establishing the optimal redox range that might help preserve sperm function and improve ART outcomes.

### 4.5 Future perspectives and potential implications of seminal redox status for porcine fertility

It is important to note that ORP measures the overall balance between oxidants and antioxidants, whereas TAC only captures a fraction of the redox homeostasis, primarily assessing the total capacity of non-enzymatic antioxidants. Future studies should therefore be directed towards evaluating the relationships between ORP and similar redox indices like the ratio between total oxidant status and TAC or between advanced oxidation protein products and TAC [55, 56]. Another key future direction for research is to establish ORP cut-off values that can differentiate between abnormal and normal seminal specimens in boars, similarly to those already established in humans (e.g., >1.34 mV/10^6^ sperm/mL [14]; >0.51 mV/10^6^ sperm/mL [18]; >1.41 mV/10^6^ sperm/mL [19]). This will require a comprehensive dataset from both fertile and infertile boars. Furthermore, this standardized approach will facilitate an in-depth exploration of other variables that may influence seminal redox dynamics, such as ejaculate fractions, breed, and seasonality. Because seminal TAC is predictive of boar fertility [42] and given the correlation found in this study between this parameter and seminal ORP, further research is warranted to determine if seminal redox status can predict boar reproductive efficiency. This could enable a simple and rapid screening of boar reproductive potential at breeding centers.

## 5. Conclusions

Our study provides evidence that ORP of ejaculate and seminal plasma were significantly correlated with the seminal TAC. Moreover, ORP of the boar ejaculate was significantly lower than that of the seminal plasma. Among sperm parameters, ORP was mainly influenced by seminal pH and sperm concentration. Therefore, sperm concentration and seminal pH must be controlled in studies assessing ORP of samples treated with oxidants/antioxidants. Interestingly, both ORP and TAC showed similar correlations with sperm parameters, being both associated with sperm concentration, mitochondrial activity, and velocity. While boar grouping on the basis of either ORP or TAC did not seem to predict sperm quality during liquid storage, seminal ORP significantly increased during the same storage time irrespective of sperm presence. The assessment of seminal ORP is a helpful and interesting tool for evaluating sperm redox homeostasis in animal andrology.

## CRediT authorship contribution statement

**Eliana Pintus:** Conceptualization, Data curation, Formal analysis, Investigation, Methodology, Project administration, Supervision, Validation, Visualization, Writing – original draft. **Maria Scaringi:** Data curation, Investigation, Methodology, Resources, Validation, Writing – review and editing. **Jef Engelen:** Data curation, Investigation, Methodology, Writing – review and editing. **José Luis Ros-Santaella:** Conceptualization, Data curation, Funding acquisition, Investigation, Methodology, Project administration, Supervision, Validation, Visualization, Writing – review and editing.

## Data availability

The data presented in this study are available on request from the corresponding authors.

## Acknowledgements

The authors gratefully acknowledge the websites NIAID NIH BioArt Source and www.flaticon.com for the illustrations used to create the graphical abstract. Full citations for the illustrations are as follows:

NIAID Visual & Medical Arts. (10/7/2024). Pig. NIAID NIH BIOART Source. bioart.niaid.nih.gov/bioart/406

NIAID Visual & Medical Arts. (10/7/2024). Mitochondria. NIAID NIH BIOART Source. bioart.niaid.nih.gov/bioart/352

NIAID Visual & Medical Arts. (10/7/2024). Human Sperm. NIAID NIH BIOART Source. bioart.niaid.nih.gov/bioart/241

www.flaticon.com. Balance free icon made by Kiranshastry. https://www.flaticon.com/free-icon/balance_1153269?related_id=1153503&origin=search

## Declaration of generative AI and AI-assisted technologies in the manuscript preparation process

During the preparation of this work the authors used Gemini to enhance the readability and language of the manuscript. After using this tool, the authors reviewed and edited the content as needed and take full responsibility for the content of the published article.

